# Human intracranial pulsatility during the cardiac cycle: a computational modelling framework

**DOI:** 10.1101/2022.05.19.492650

**Authors:** Marius Causemann, Vegard Vinje, Marie E. Rognes

## Abstract

**Background:** Today’s availability of medical imaging and computational resources set the scene for high-fidelity computational modelling of brain biomechanics. The brain and its environment feature a dynamic and complex interplay between the tissue, blood, cerebrospinal fluid (CSF) and interstitial fluid (ISF). Here, we design a computational platform for modelling and simulation of intracranial dynamics, and assess the models’ validity in terms of clinically relevant indicators of brain pulsatility.

**Methods:** We develop finite element models of fully coupled cardiac-induced pulsatile CSF flow and tissue motion in the human brain environment. The three-dimensional model geometry is derived from magnetic resonance images (MRI), features a high level of detail including the brain tissue, the ventricular system, and the cranial subarachnoid space (SAS). We model the brain parenchyma at the organ-scale as an elastic medium permeated by an extracellular fluid network and describe flow of CSF in the SAS and ventricles as viscous fluid movement. Representing vascular expansion during the cardiac cycle, a pulsatile net blood flow distributed over the brain parenchyma acts as the driver of motion. Additionally, we investigate the effect of model variations on a set of clinically relevant quantities of interest.

**Results:** Our model predicts a complex interplay between the CSF-filled spaces and poroelastic parenchyma in terms of ICP, CSF flow, and parenchymal displacements. Variations in the ICP are dominated by their temporal amplitude, but with small spatial variations in both the CSF-filled spaces and the parenchyma. Induced by ICP differences, we find substantial ventricular and cranial-spinal CSF flow, some flow in the cranial SAS, and small pulsatile ISF velocities in the brain parenchyma. Moreover, the model predicts a funnel-shaped deformation of parenchymal tissue in dorsal direction at the beginning of the cardiac cycle.

**Conclusions:** Our model accurately depicts the complex interplay of ICP, CSF flow and brain tissue movement and is well-aligned with clinical observations. It offers a qualitative and quantitative platform for detailed investigation of coupled intracranial dynamics and interplay, both under physiological and pathophysiological conditions.

## Introduction

The pulsating brain environment features a unique and dynamic interplay between blood influx and efflux, cerebrospinal fluid (CSF) flow in and between the cranial and spinal compartment, intracranial pressures (ICPs), brain tissue movement and interstitial fluid (ISF) flow. Alterations in the dynamics of ICP or CSF flow are associated with central nervous system disorders [60] such as hydrocephalus [32, 44], Alzheimer’s disease and multiple sclerosis [47]. Moreover, better understanding of CSF flow characteristics could play an important role for targeted drug delivery [41]. Progress in magnetic resonance imaging (MRI) has allowed for non-invasive measurements of CSF flow, blood flow, and brain tissue deformation [4, 48]. Over the last decade, computational modelling of brain mechanics have emerged as a promising complementary tool to obtain high fidelity and high resolution models and predictions of intracranial dynamics [35].

Computational studies of intracranial pulsatility have mainly focused on either the brain parenchyma [26, 25, 58] or the flow of CSF through the ventricular system and the spinal and cerebral subarachnoid spaces (SAS) [33, 29, 53, 59]. Such *decoupled* approaches do not fully account for the close interactions between the brain tissue and the surrounding CSF, and the potential exchange between CSF and ISF. In contrast, *coupled* fluid-structure interaction models allow for simultaneous computation of flow and pressure in the CSF-spaces as well as the solid displacement and stresses in the brain parenchyma. Linninger et al. [34] proposed a model of CSF flow in the SAS and ventricles coupled with porous media flow through the brain parenchyma driven by an oscillatory inflow boundary condition at the choroid plexus. Sweetman et al. [51] introduced a 3D model of CSF flow with fluid-structure interaction driven by a moving lateral ventricle wall. Tully and Ventikos [54] investigated the coupling of poroelasticity and free fluid flow using an idealized brain model. Gholampour [23] used a coupled model of CSF flow and brain viscoelasticity – again driven by a CSF source in the lateral ventricles to compare flow patterns in healthy and hydrocephalic subjects.

Based on multi-modal MR imaging, Balédent [4] proposed that cardiac cycle-induced intracranial pulsatility is driven by the following sequence of events. During systole, arterial blood flow into the brain exceeds the venous outflow, the brain expands, ICP increases, and CSF is displaced into the spinal canal. Subsequently, during diastole, venous outflow dominates the vascular dynamics, leading to a decrease of ICP and a reversal of CSF flow. A key question is whether and to what extent computational models can integrate this view of intracranial dynamics, driven by the cardiac-induced expansion of blood vessels in the brain tissue [4], with clinical observations of ICP [19], ICP differences [20, 59], and CSF flow.

In this paper, we therefore propose a computational model of intracranial dynamics coupling the pulsatile motion of CSF, brain tissue and ISF during the cardiac cycle. We represent the brain parenchyma at the organ-scale as an elastic medium permeated by an extracellular network saturated by CSF/ISF. Flow of CSF in the SAS and ventricles is modelled as a viscous fluid under low Reynolds numbers i.e. via the Stokes equations. Crucially, we employ a pulsatile net blood flow distributed over the brain parenchyma as the driver of motion. This fully coupled computational model enables studies of the entire intracranial system dynamics. Specifically, the model predicts the brain displacement, intracranial pressures within the parenchyma, in the SAS, and in the ventricular system, and CSF and ISF flows. Several model variations (e.g. parameter regimes) were also tested to assess the sensitivity to different parameters. Overall, our computational results agree well with clinical observations of ICP, stroke volumes, and brain displacements, and thus introduces a promising computational approach to study intracranial pulsatility driven by intraparenchymal blood flow.

## Methods

### Domains and boundaries

We represent the brain parenchyma as a three-dimensional domain Ω_*p*_, and the surrounding CSF-filled spaces by Ω _*f*_ (Figure 1a). These two domains share a common boundary Σ = Ω _*f*_ *∩*Ω_*p*_ with normal vector **n**, pointing from Ω _*f*_ to Ω_*p*_ on Σ and outwards on the boundary *∂* Ω. Further, Γ_skull_ denotes the outer boundary of the CSF space where the rigid skull encloses the cranial cavity (Figure 1a). The lower boundary of the domain (at the C3 level) is split into two segments: the caudal continuation of the spinal cord is labeled Γ_*SC*_, while Γ_*SAS*_ describes the boundary to the spinal SAS. To obtain a computational mesh of these domains, we manually segmented the *full head MRI scan* data set provided by Slicer3D [22, 30], and extracted the constituents of the ventricular system, the cranial SAS and the brain parenchyma (Figure 1c). The surfaces of the segmented regions were meshed using the Surface Volume Meshing Toolkit (SVMTK) [57]. The volumes and diameters of the relevant mesh substructures, as listed in Table 1, are within clinically reported ranges. The computational mesh consists of 4526016 mesh cells, 796303 mesh vertices and a maximal (minimal) cell diameter of 6.7 mm (0.2 mm).

**Table 1.** Comparison of the generated computational head model and experimentally determined values in healthy subjects with respect to the dimensions of the brain’s substructures; C3 is the third cervical vetrebra level of the spine

| Substructure dimension | This study | Literature value | Reference |
| --- | --- | --- | --- |
| Lateral ventricles volume | 24.01 ml | 9.82 ml (normal)<br>250.2 ml (hydrocephalic) | Linninger et al. [34] |
| Third ventricle volume | 3.60 ml | 2.48 ml | Linninger et al. [34] |
| Fourth ventricle volume | 2.69 ml | 3.31 ml | Linninger et al. [34] |
| Subarachnoid space volume | 292.08 ml | 179 ml (only cranial) | Chazen et al. [14] |
| Aqueduct diameter | 2.88 mm | 1.5 to 3.0 mm | Haines and Mihailoff [27] |
| Spinal canal (C3) diameter | 12 mm | 9.4 to 17.2 mm | Ulbrich et al. [55] |
| Spinal coord (C3) diameter | 7 mm | 6.0 to 9.6 mm | Ulbrich et al. [55] |
| Brain parenchyma total volume | 1369.54 ml | 1130 ml (women)<br>1260 ml (men) | Cosgrove, Mazure, and Staley [16] |

**Figure 1.**
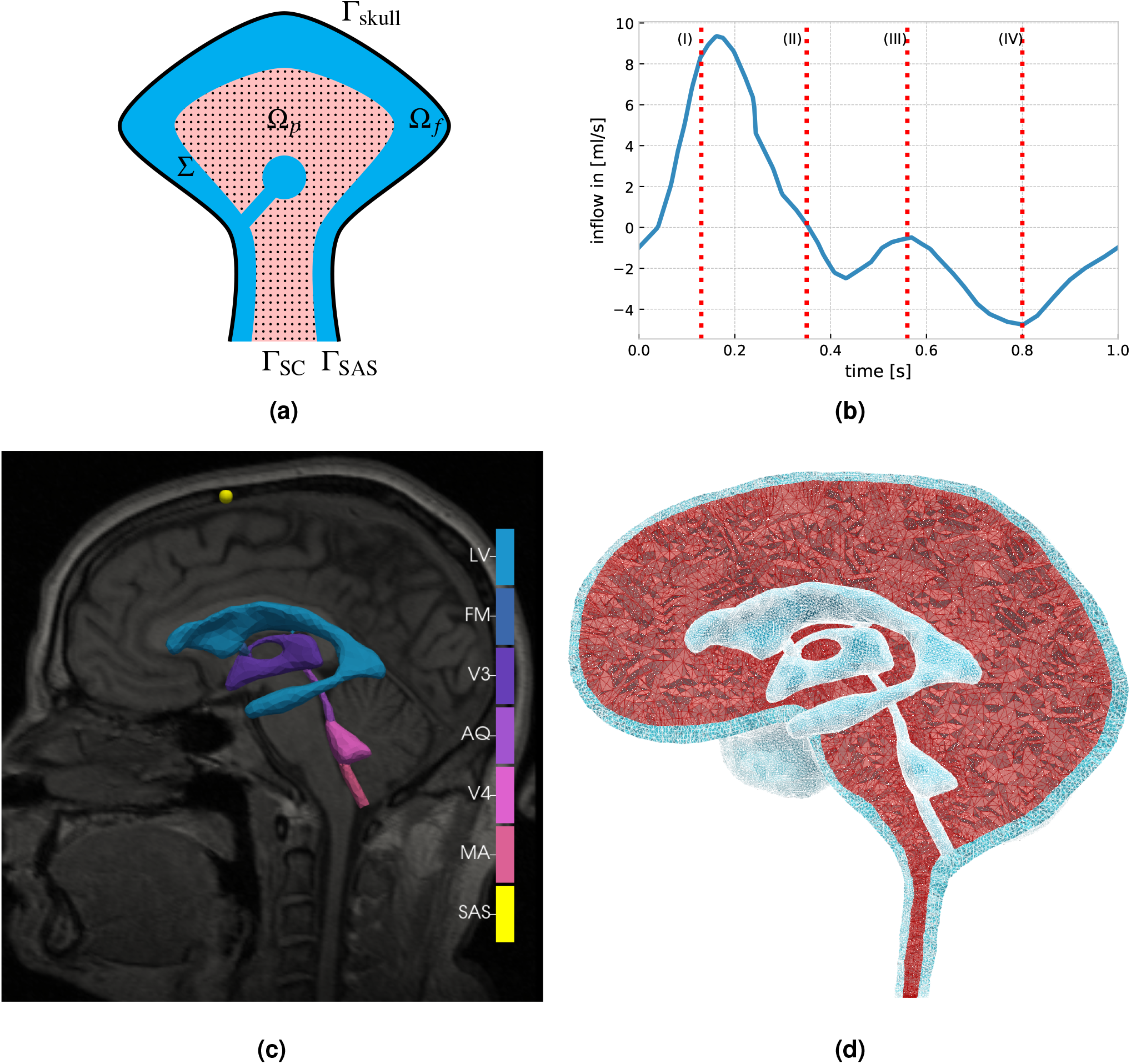
**a)** Sketch of the domains representing the brain parenchyma (Ω_*p*_, pink) and the CSF-filled spaces (Ω _*f*_, blue). The interface of both domains is denoted by Σ. Additionally, the boundaries Γ_skull_ at the skull, Γ_SC_ at the spinal cord and Γ_SAS_ at the spinal SAS are highlighted; **b)** Net blood inflow during the cardiac cycle with four different phases: (I) early systole - high net blood inflow; (II) end of net blood inflow phase; (III) brain equilibrium phase (arterial inflow and venous outflow almost match); (IV) high net outflow of blood (data extracted from Balédent [4]); **c)** The MRI image used for the mesh generation and the segmented parts of the ventricular system: LV - lateral ventricles, FM - foramina of Monro, V3 - third ventricle, AQ - aqueduct of Sylvius, V4 - fourth ventricle, MA - median aperture, SAS - probe point in the subarachnoid space; **d)** sagittal view of the mesh, displaying the ventricular system, cranial SAS (both light blue) brain parenchyma (red).

### Governing equations

#### The brain parenchyma

We regard brain tissue as a linear poroelastic medium permeated by a single fluid network representing an extracellular CSF/ISF-space. The equations of linear poroelasticity express conservation of momentum for the solid elastic matrix and the mass conservation of a diffusive flow within the medium [9]. Due to its robustness in case of materials close to the incompressible limit or with low storage capacity, we chose a three-field formulation, based on the displacement **d**, fluid (pore) pressure *p*_*p*_ and the additional total pressure *ϕ*, which is defined as *ϕ* = *αp*_*p*_ *−λ* div **d** [31, 39]. With the infinitesimal strain tensor 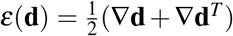 and a volume source term *g*, the equations read as follows:

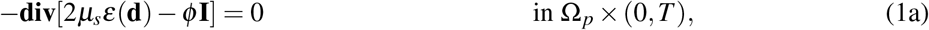

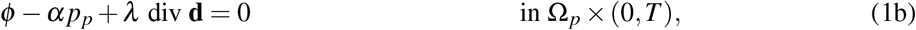

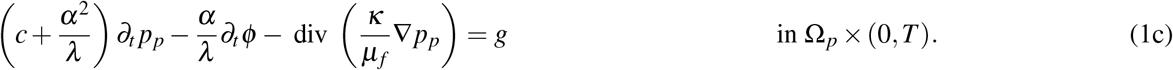

Here, *κ* represents the permeability, *c* the specific storage coefficient, and *α* the Biot-Willis coefficient. The identity operator is **I**. The linear isotropic solid matrix is parameterized with the Lamé constants *µ*_*s*_ and *λ*, while the fluid permeating the pores has viscosity *µ*_*f*_.

#### CSF compartments

We model the flow of CSF in the ventricular system and SAS by the time-dependent Stokes equations for the CSF velocity *u* _*f*_ and fluid pressure *p* _*f*_. The Stokes equations represent flow under low Reynolds numbers typically observed in the CSF compartments; Howden et al. [29] report an average Reynolds number of *Re*_*av*_ = 0.39 with a maximum value of *Re*_*max*_ = 15 in the CSF-filled spaces of the cranium during the cardiac cycle. Under these assumptions, the equations reads as follows:

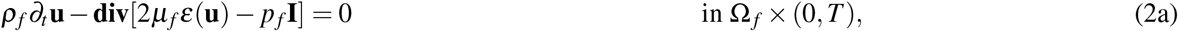

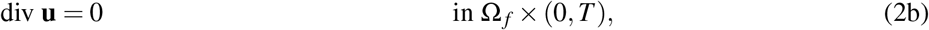

with the strain rate tensor 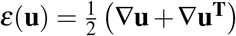, constant CSF density *ρ*_*f*_, and constant CSF viscosity *µ*_*f*_.

### Net blood flow as a driver of pulsatility

We induce motion in the system via a vascular expansion through net flow of blood into the brain parenchyma, modelled by a pulsatile source term *g* in (1c). We define net blood flow as the difference between arterial blood inflow and venous blood outflow over time. As Biot’s equations include only one fluid network, we treat the net blood flow as a source term in this single fluid compartment. This simplification can be justified by the similarity of the effect of an inflow of blood and/or ISF: both lead to a volumetric expansion of the brain parenchyma and an increase of fluid pressure. We let *g* vary in time, but be spatially uniform, and employ a measured net blood inflow time series from Balédent [4] (Figure 1b). The rapid inflow of arterial blood during early systole (phase I) increases the cranial blood volume, until venous outflow balances the arterial inflow, ending the net inflow of blood (phase II). Next, after a brief equilibrium (phase III), the venous outflow exceeds the arterial inflow (phase IV) and sets the cerebral blood circulation up for the next cycle.

### Transmission, boundary and initial conditions

We augment the above governing equations by the following transmission (interface), boundary and initial conditions.

#### Transmission Conditions

Based on first principles, we require the following equations to hold on the interface Σ between the porous and the fluid domain:

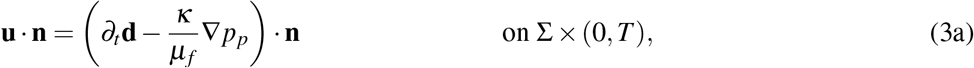

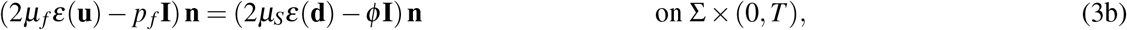

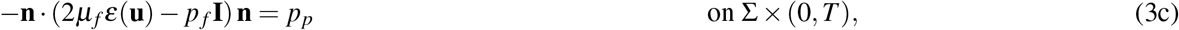

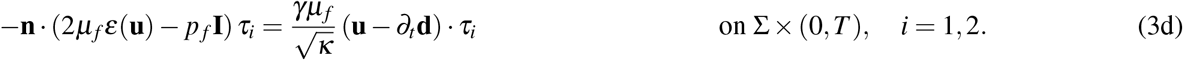

Here, to complement the normal **n**, *τ*_*i*_ (*i* = 1, 2) we define orthogonal tangent vectors to the interface, and *γ >* 0 is the slip rate coefficient, which is a dimensionless constant depending only on the structure of the porous medium. Here, (3a) enforces continuity of the normal flux on the interface, (3b) conserves momentum, while (3c) accounts for the balance of total normal stress. The last interface condition (3d) is the Beavers-Joseph-Saffman (BJS) condition, which states that the jump in the tangential velocities across the interface is proportional to the shear stress on the free flow side of the interface [7, 46, 37].

#### Boundary Conditions

Assuming a rigid skull, we set no-slip conditions on the skull boundary Γ_skull_:

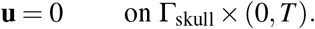

For the spinal cord boundary Γ_SC_, we assume no displacement and no flux:

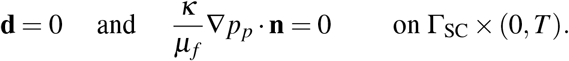

To represent the compliance of the spinal compartment, we assume an exponential relationship between ICP and additional volume [36, 61, 52]:

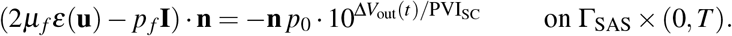

The pressure-volume index (PVI_SC_) represents a clinical measure of the compliance of the spinal compartment, *p*_0_ is the initial pressure of the system and Δ*V*_out_(*t*) is the total additional volume of CSF in the spinal compartment. The latter equals the volume of CSF that has left the domain over the corresponding part of the boundary Γ_SAS_, and is calculated as follows:

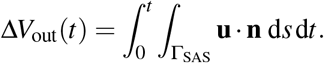

This allows for the pulsatile motion of CSF in and out of the domain.

#### Initial Conditions

Finally, we assume that the system is initially at rest with an initial pore pressure *p*_0_:

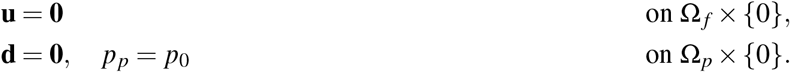

### Material parameters

Material parameters were selected based on literature values and are summarized in Table 2.

**Table 2.**
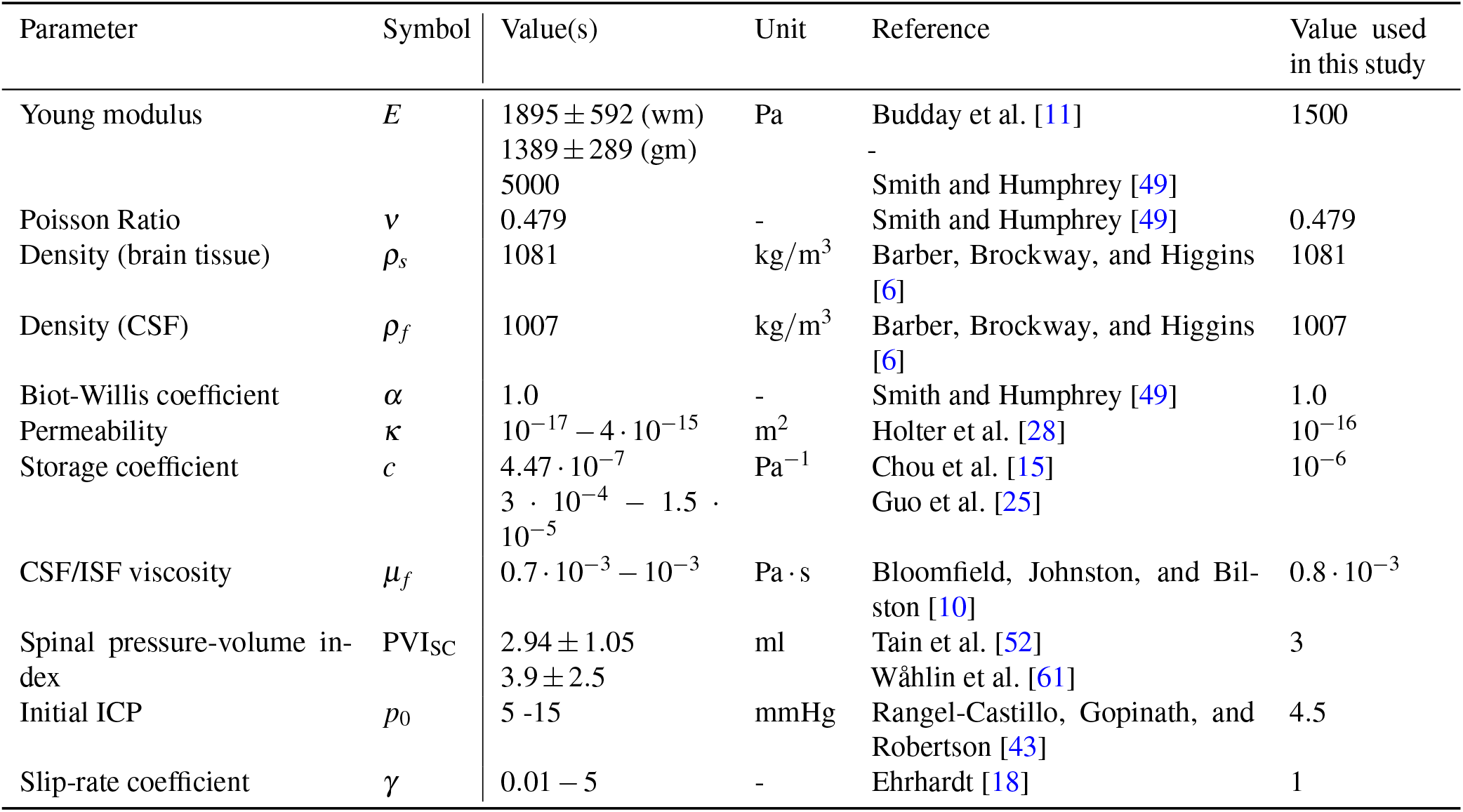
Summary of material parameters, including references to values from previous studies.

#### Quantities of interest

Primary clinical quantities of interest are the ICP and CSF flow rates and volumes in the foramen magnum or across the aqueduct [60]. In our computational model, we identify the ICP as the (fluid) pressure *p* _*f*_ in the CSF compartment(s) and as the total pressure *ϕ* in the parenchyma, which incorporates both the pore pressure and the stress exercised by the elastic matrix. We place virtual/computational pressure probe points inside the lateral ventricles, in the cranial SAS at the upper convexity of the skull, and inside the fourth ventricle (Figure 1c). Flow rates within the ventricular system and into the spinal compartment are obtained by spatial integration of the computed CSF flow across boundaries between the different parts of the ventricular system or across the spinal external boundary, respectively. Specifically, we define the following set of quantities of interest:

i. the peak volumetric flow rate in the aqueduct,
ii. the aqueduct stroke volume, corresponding to the net volume of fluid pulsating back and forth in the aqueduct over the cardiac cycle (maximum of the cumulative flow volume),
iii. the peak tissue displacement,
iv. the (peak) transmantle pressure gradient, computed as the (peak) pressure difference between the virtual probe points in the cranial SAS and the lateral ventricles and divided by the distance between these points,
v. the temporal nadir-to-peak (i.e, diastolic to systolic) amplitude of pressure in the lateral ventricles,
vi. the spinal stroke volume, corresponding to the net volume of fluid pulsating back and forth into the spinal compartment over the cardiac cycle.

Results are reported from the last of three cardiac cycles to limit the influence of the initial data.

#### Model variations

The effect of the model’s parameterization is of particular interest due to the uncertainty of the chosen parameters. Additionally, variations of material parameters offer insights into the relation of changing material characteristics (possibly caused by diseases or ageing) and alterations in the pulsatile motion of the brain. Since an extensive exploration of the parameter space of the model is out of scope for this work, we restrict our analysis to a collection of selected deviations from the standard model (Table 3). For model A, we increase the pressure-volume index *PVI* = 10 ml, which corresponds to a larger spinal compliance. Model B represents stiffer brain parenchyma (Young Modulus *E* = 3000 Pa) while in model C we increase the compressibility of the brain (Poisson ratio *ν* = 0.4). Finally, model D features a greater storage coefficient (*c* = 10^*−*5^ Pa^*−*1^), which reduces the rise of pressure with additional fluid volume inside the poroelastic parenchyma and hence models larger intracranial compliance.

**Table 3.**
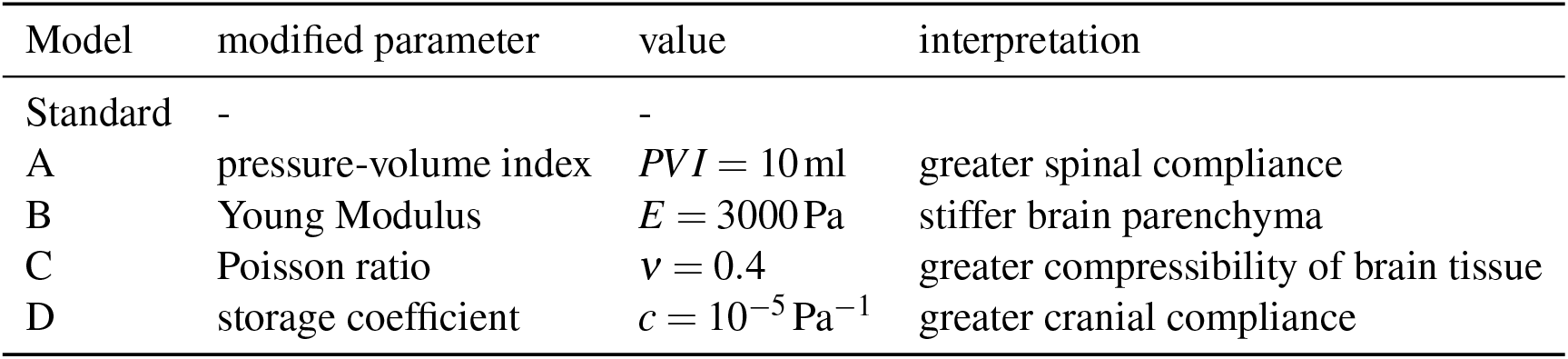
Overview of the selected models, their deviation from the standard parameterization and the corresponding interpretation

#### Numerical methods & software

The complete system was solved via a fully coupled strategy with a an implicit Euler finite difference discretization in time and a finite element method in space, following [45]. We approximate the vector-valued unknowns, i.e. the tissue displacement and fluid velocity, with continuous piecewise quadratic polynomials, while continuous piecewise linear functions are employed for the pore pressure, total pressure, and fluid pressure. The model is implemented with the finite element software *FEniCS* [1] and its extension to multiphysics problems *multiphenics* [5]. The resulting linear system is factorized and solved in every time step with the direct solver *MUMPS* [2, 3], employing a hybrid approach of distributed and shared memory parallelism (via OpenMP and MPI).

We performed convergence tests against smooth manufactured solutions to verify the accuracy of the discretization and further verified the computations using mesh and time step convergence tests (Supplementary Figure 9 and Figure 10).

## Results

The cardiac-induced influx of blood to the brain parenchyma induces a complex interplay between the CSF-filled spaces and poroelastic parenchyma in terms of intracranial pressures and pressure gradients, CSF and ISF flow, and parenchymal displacements.

### Intracranial pressure

At the beginning of the cardiac cycle, the ICP rapidly and nearly uniformly rises from its initial value of 4.5 mmHg to reach a peak of 8.4 mmHg after approximately 0.3 s (Figure 2). Subsequently, it steadily decreases until the initial value is reached again and the next cycle begins. The nadir-to-peak pressure variation in time is close to 4.0 mmHg, whereas the spatial differences are several orders of magnitude smaller. The transmantle pressure gradient between the lateral ventricles and upper convexity of the SAS peaks at 0.18 mmHg/m (Figure 2b). The maximal gradient between the lateral and the fourth ventricle is almost three times larger, reaching a peak value of 0.41 mmHg/m. The temporal variations in these pressure gradients oscillate with higher frequency than the cardiac cycle.

**Figure 2.**
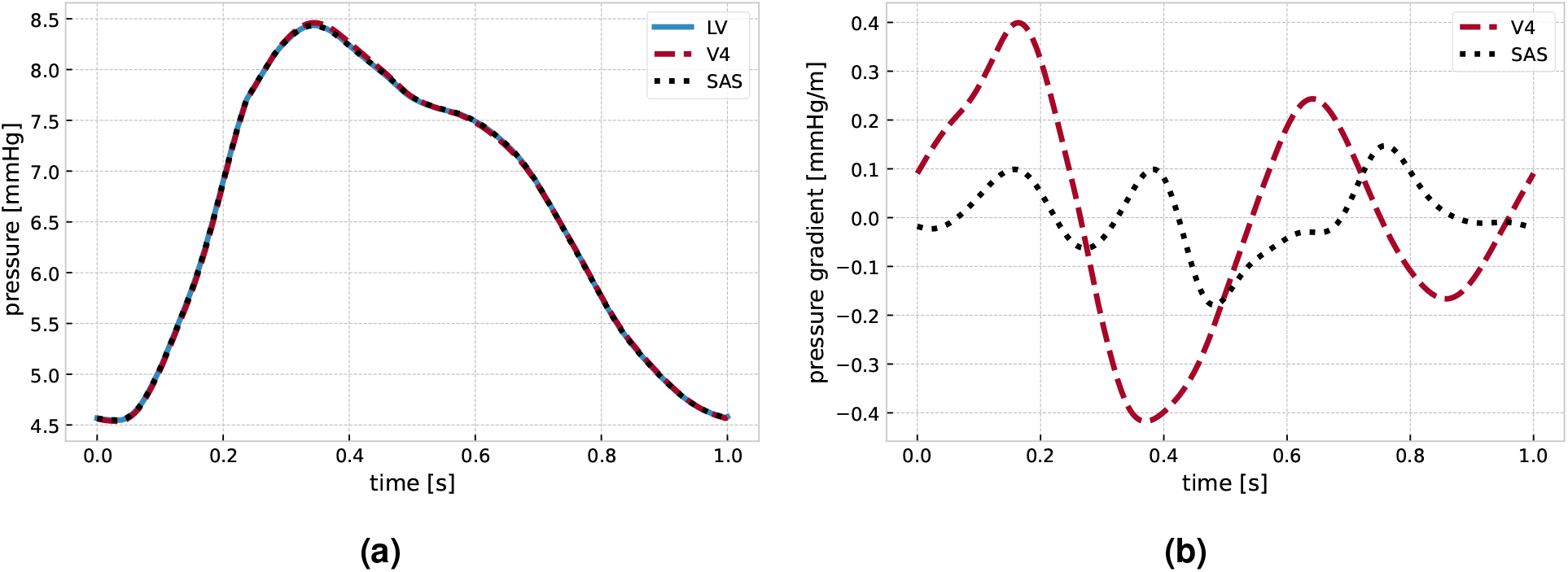
(a) Evolution of the ICP inside the lateral ventricles, in the cranial SAS at the upper convexity of the skull, and inside the fourth ventricle. (b) Intracranial pressure gradient from the lateral ventricles to the upper convexity of the cranial SAS (black) and the fourth ventricle (red).

The spatial ICP distribution differs between the four phases of the cardiac cycle (see Figure 3 sagittal, coronal and transversal views). In phase I (early systole), we observe the largest spatial pressure variation of the four phases.

**Figure 3.**
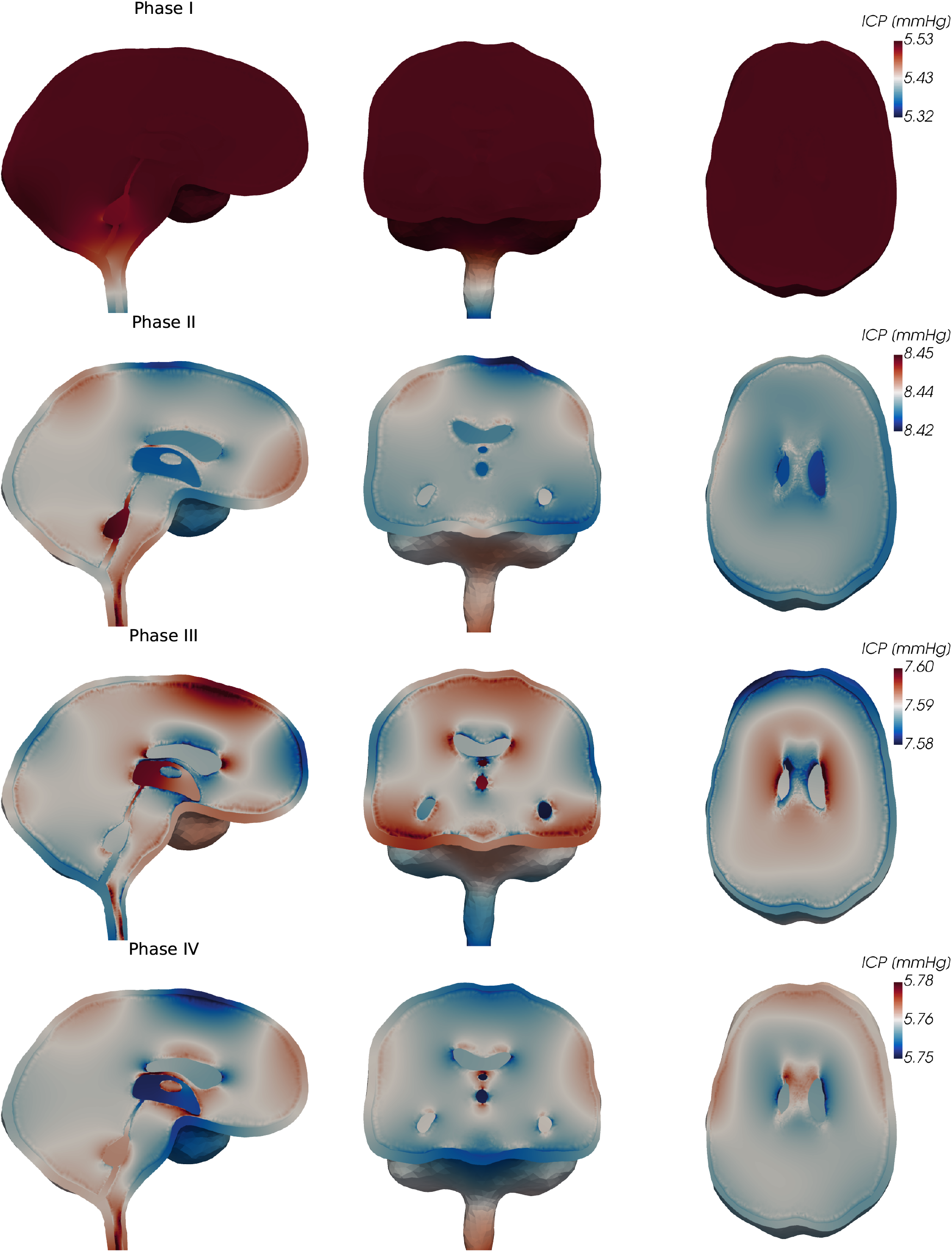
Sagittal, coronal, and transversal views of the ICP (fluid pressure in the CSF-filled spaces and total pressure in the parenchyma) during phases I–IV of the cardiac cycle. Note that the color scale changes between the different phases (rows).

While the ICP in the parenchyma, the ventricular system, and the cranial SAS are nearly equal, ICP decreases in the dorsal direction from the craniospinal junction at the foramen magnum. This results in a pressure drop of 0.21 mmHg from the cranium to the spinal compartment. Additionally, we observe a slightly lower pressure in the fourth ventricle compared to the third ventricle and surrounding tissue. In phase II (end of net blood inflow), spatial ICP differences amount to 0.03 mmHg, less than 15% of that of phase I. The peak pressure is now observed at the lowest point of the cervical spine and in the fourth ventricle. The pressure differences in the ventricular system thus reverse: highest values occur in the fourth ventricle, decreasing towards the third ventricle and resulting in a small pressure gradient over the aqueduct. Next, phase III (brain equilibrium) is characterized by small spatial pressure differences of less than 0.02 mmHg. Inside the ventricular system, the pressure difference over the aqueduct once again reverses, and the largest pressure is obtained in the third ventricle. Finally, in phase IV (high net blood outflow), the pressure increases from the craniocervical junction in the caudal direction. The lowest pressure occurs at the frontal part of the upper convexity of the skull and in the third ventricle. The pressure difference across the aqueduct reverses yet again.

### CSF flow patterns

The differences in pressure distributions induce characteristically different CSF flow patterns across the cardiac phases (Figure 4). In phase I, CSF rushes out of the cranium into the spinal canal reaching a peak velocity of 78.5 mm/s at the craniocervical junction. Simultaneously, a slower, caudally-directed flow of CSF occurs within the cranial SAS at velocity magnitudes on the order of 10 mm/s. CSF inside the ventricular system is displaced downwards through the fourth ventricle and the median aperture. During phase 2, CSF flows from the lateral ventricles through the foramina of Monro into the third ventricle. Flow in the aqueduct is nearly stagnant, while flow in the median aperture reverses and is directed into the fourth ventricle. Simultaneously, the caudal CSF flow in the upper convexity of the cranium and the outflow into the spinal compartment continue on a smaller scale. In phase 3, almost no flow occurs into the spinal compartment. Inside the ventricular system, we again observe a reversal of flow directions: CSF moves in the median aperture in the dorsal direction and runs in the opposite direction at the level of the aqueduct and third ventricle. Finally, in phase 4, we observe the return of CSF from the spinal compartment into the cranium. CSF flows through the spinal canal, the cranial SAS, and the lower part of the ventricular system and thereby completes its cycle.

**Figure 4.**
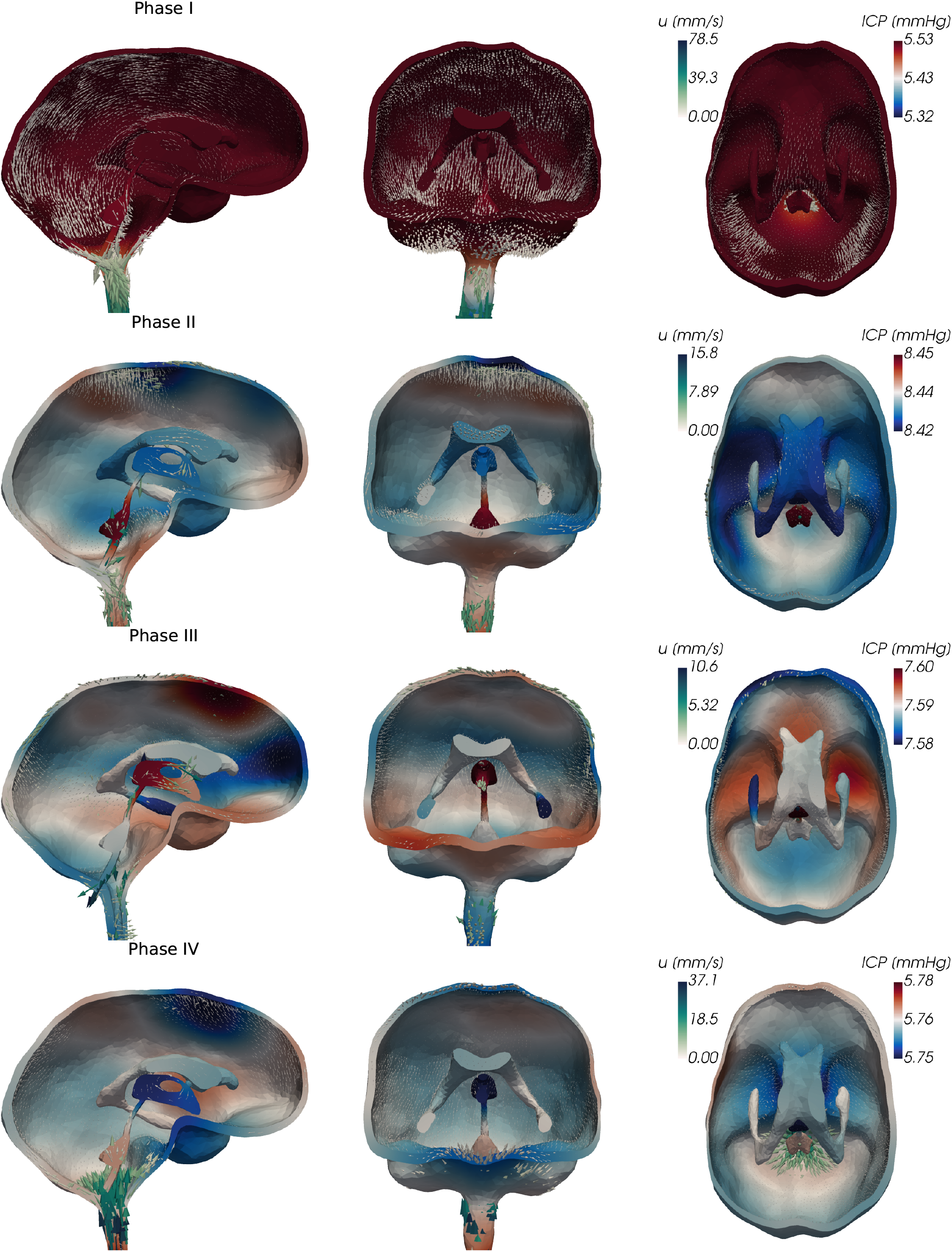
Sagittal, coronal, and tranverse views of the pressure (ICP) and fluid velocity **u** in the CSF-filled spaces of the cranium during different phases of the cardiac cycle. (Logarithmic scaling of the arrows representing the velocity.)

In addition to this global description of CSF flow, we consider the flow rates and volumes in the ventricular system and at the cervical level in more detail (Figure 5). The largest flow rate occurs into the spinal canal, where up to 6 ml/s leave the cranium into the spinal compartment (Figure 5a). This CSF-spinal flow rate thus corresponds to approximately two-thirds of the amplitude of the net blood inflow. The resulting stroke volume is 0.8 ml and corresponds to the peak value of the spinal cumulative flow volume at 35% of the cardiac cycle (Figure 5b). The ventricular flow rates are at least one order of magnitude lower than those of the spinal canal, reaching at most 0.22 ml/s at the transition from the fourth ventricle to the median aperture. In the aqueduct, we observe a peak flow rate of 0.07 ml/s and a stroke volume of 0.013 ml (Figure 5c, Figure 5d). Notably, within each cardiac cycle, the flow reverses its direction multiple times. In the lower parts of the ventricular system (median aperture, aqueduct), flow initially takes place in the dorsal direction and changes its direction three times. At the level of the foramina of Monro, we observe a short phase of flow into the lateral ventricles at the beginning of the cycle and again three reversals of direction. Thus, the time of the flow rate peaks in the upper regions of the ventricular system are delayed compared to the lower regions (Figure 5c)

**Figure 5.**
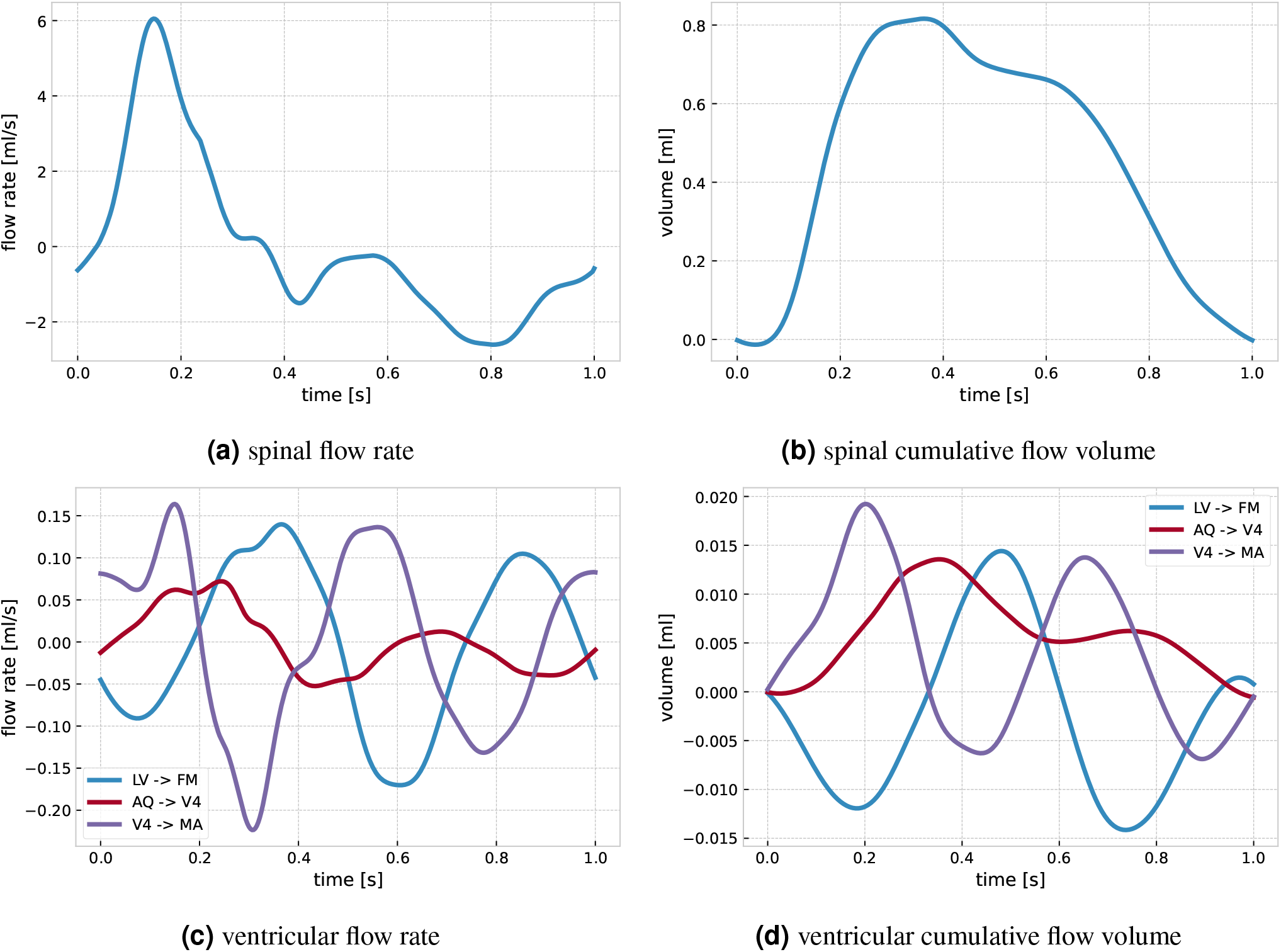
Volumetric flow rates and stroke volumes within the ventricular system and into the spinal compartment. LV -> FM denotes flow from the lateral ventricles into the foramina of Monro, AQ -> V4 from the aqueduct into the fourth ventricle, and V4 -> MA from the aqueduct into the median aperture (cf. Figure 1c).

Interstitial flow velocities and volumes within the parenchymal tissue pulsate with the cardiac cycle but are generally small (peak velocity magnitude less than 1.9 *µ*m/s, and peak spatial average of 0.13 nm/s). The exchange between ISF and CSF is on the order of nanoliters per second which is negligible compared to flow rates in the spinal canal (on the order of ml/s).

#### Brain parenchyma displacements

During early systole (phase I), a large dorsal deformation occurs, especially of the infratentorial part of the brain (Figure 6). A peak displacement magnitude of 0.22 mm is found in the brain stem 12 % into the cardiac cycle. After 35% of the cycle (in phase II), most of the infratentorial brain regions have return to their original configuration. In this phase, the displacement predominately occurs at the anterior and posterior ends, and we observe a rotational movement of the brain around its center. While the posterior regions are deformed downwards, the frontal region moves up and backwards. In the third phase, the overall pattern changes only slightly. Specifically, the anterior displacement decreases and the center of rotation moves forward. In the final phase of the cardiac cycle, the displacement magnitude decreases substantially and the remaining displacement is predominantly in the frontal superior parts in an upwards direction and in the central inferior region of the brain in the caudal direction. Throughout the cycle, we note some radial displacements of the spinal cord.

**Figure 6.**
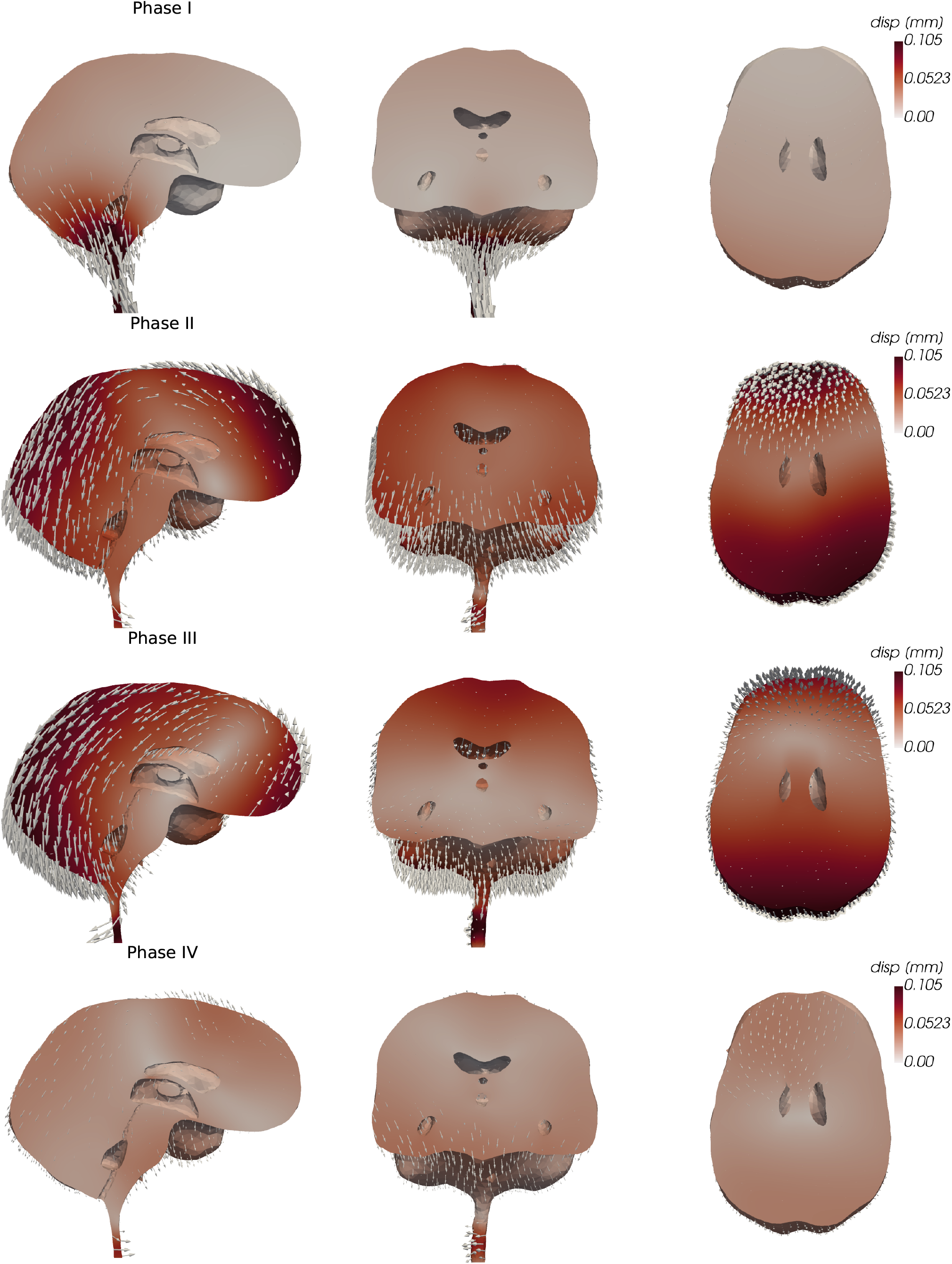
Sagittal, coronal, and transverse views of the brain parenchymal displacement during different phases of the cardiac cycle. The glyph arrows representing the displacement are amplified by a factor of 200.

#### Role of brain and spinal cord compliances

The set of quantities of interest predicted by the different computational models (models A–D) differ from the standard model (Figure 7). For all quantities of interest, the outputs of the models range between 19 % and 166 % relative to the standard model.

**Figure 7.**
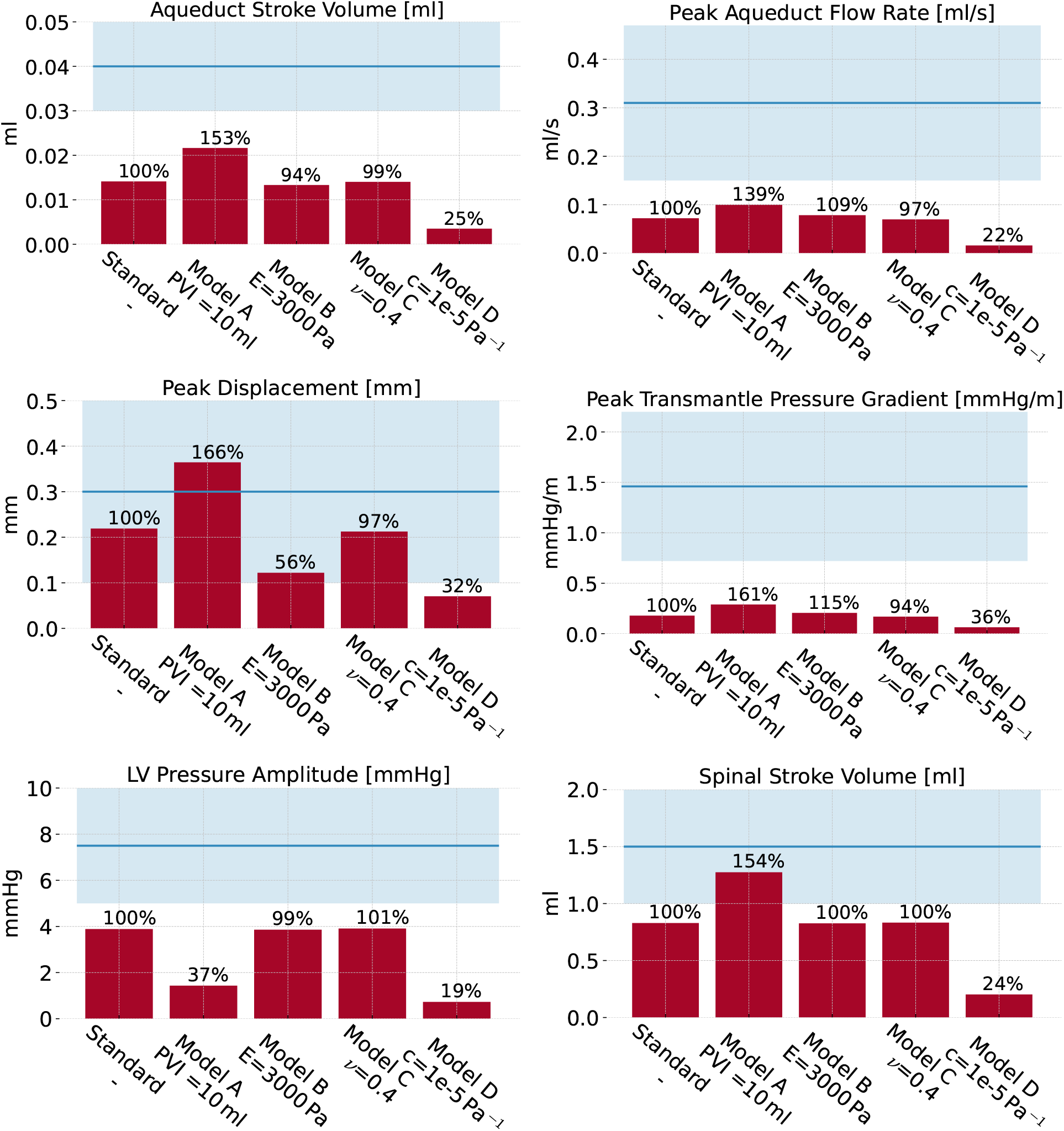
Overview of clinical quantities of interest of a set of model variations from the standard parameterization. The blue horizontal bar represents the range of physiologically realistic values with the blue line indicating the mean value.

#### Increased spinal compliance

Increasing the spinal compliance by increasing the spinal pressure-volume index (Model A), yields increased aqueduct stroke volumes, spinal stroke volumes and peak aqueduct flow rates (by 53%, 54%, 39% respectively) relative to the standard model. In addition, the peak displacement is increased by 66%, and the peak transmantle pressure gradient increases by 61%. Conversely, the total pressure variation in the lateral ventricles is substantially reduced, by 63%. In addition, the ICP curve changes characteristics (Supplementary Figure 11). With increased spinal compliance, additional peaks (P1, P2, P3) are seen in the ICP signal.

#### Increased brain stiffness

Increasing the brain stiffness (Model B) reduces the peak brain displacement by 44%. The other clinical quantities of interest remain unchanged.

#### Increased brain compressibility

Increasing the brain compressibility (Model C) yields only negligible changes in clinical quantities of interest. The largest difference relative to the standard model is observed for the peak transmantle pressure gradient, and only amounts to a 6% decrease.

#### Increased storage coefficient

Decreasing the brain parenchyma’s poroelastic storage coefficient (Model D) results in substantial decreases in the set of clinical quantities of interest computed. The aqueduct stroke volume, spinal stroke volume and peak aqueduct flow rates are reduced by by 75%, 76%, and 78%, respectively. The peak displacement is decreased by 68%, and the peak transmantle pressure gradient by 64%. Similarly, the total pressure variation in the lateral ventricles is reduced by 81%.

## Discussion

We have presented a three-dimensional computational model of fully coupled cardiac-induced pulsatile CSF flow and tissue motion in the human brain environment. Variations in the ICP were dominated by their temporal amplitude, but with small spatial variations in both the CSF-filled spaces and the parenchyma. The ICP variations induce substantial ventricular and cranial-spinal CSF flow, some flow in the cranial SAS, and small pulsatile ISF velocities in the brain parenchyma. Investigating the displacement of parenchymal tissue, we found a funnel-shaped deformation in dorsal direction at the beginning of the cardiac cycle, followed by a rotational motion around an axis normal to the brain’s sagittal plane. Moderate variations in the brain and spinal cord compliances altered model outputs.

The temporal pressure variations are in good agreement with previous clinical reports. Wagshul, Eide, and Madsen [60] reported typical nadir-to-peak ICP amplitudes of 5 to 10 mmHg for healthy subjects, which is only slightly higher than the 4 mmHg obtained here. Considering the morphology of the ICP waveform, notable differences between individuals seem to exist: while the general cardiac cycle pattern of increasing and decreasing pressure persists across subjects, many clinical studies have reported several peaks in the ICP signal (P1, P2, P3) [60, 56]. Unnerbäck, Ottesen, and Reinstrup [56] suggested that the first peak (P1) is caused by the rapid rise of blood inflow, while the following peaks may be related to subsequent resonance phenomena. Carrera et al. [12] related P1 to peak arterial inflow, while P2 and P3 were related to peak values in cerebral arterial blood volume. In our (standard) computational model, the peak in ICP signal is related to the change of sign in the net blood flow curve. However, additional peaks (P1, P2, P3) occur when the spinal compliance is increased. We note that our computed ICP curve lies well within the range of clinically reported curves by Ziółkowski et al. [62] and closely resembles the in-vitro modelling results by Benninghaus et al. [8]. A transmantle pressure gradient is hypothesized to drive the development of hydrocephalus [50, 19], though with recent findings also pointing at genetic factors [17]. Stephensen, Tisell, and Wikkelsö [50] reported no static transmantle pressure gradient, which agrees with the small pulsatile pressure gradients (peaking at 0.06 *−* 0.30 mmHg/m) predicted here. Taking the pulsatile nature of the ICP into account, Eide [19] measured higher amplitudes in the lateral ventricles compared to the parenchymal tissue close to the skull. Similarly, Vinje et al. [59] found pulsatile ICP gradients with average amplitudes of 1.46 *±* 0.74 mmHg/m, which is roughly one order of magnitude higher than the pulsatile transmantle gradient obtained in this work. Complementary to these clinical findings in (suspected) hydrocephalic patients, Linninger et al. [34] used computational fluid dynamics to compute maximal transmantle pressure differences of 10 Pa in healthy, and 30 Pa in hydrocephalus patients. In a subsequent modeling paper Sweetman et al. [51] predicted a maximal transmantle pressure difference in healthy individuals of 4 Pa. Assuming a distance of 6 cm between the lateral ventricles and the SAS, these pressure differences correspond to pressure gradients of approximately 0.5 *− −* 1.25 mmHg/m for healthy individuals and 3.75 mmHg/m for hydrocephalus patients. The computed transmantle pressure gradient is likely influenced by a number of model choices including the geometry representation, material parameters and importantly the assumed homogeneous net blood flow.

Consistent with the comparatively small spatial pressure differences computed, we find flow rates and stroke volumes in the ventricular system at the lower range of previous reports. The peak aqueductal flow rate and the spinal stroke volume of our standard model reach 70 % and 80 %, respectively, of the values reported by Wagshul, Eide, and Madsen [60]. However, with a higher spinal compliance, the computed spinal stroke volume (1.25 ml) is within the clinical range. This finding represents a different distribution of compliance in the overall system: a higher spinal compliance allows more CSF to leave the cranium into the spinal compartment. Furthermore, our computed aqueduct stroke volume (13 *µ*l) is lower than measured values of 30 to 50 *µ*l [60]. Balédent [4] suggested that the contribution of the ventricular system to the regulation of ICP is low compared to the effect of cervical CSF outflow. This conforms with our results since the aqueductal flow peaks later than the cervical outflow and reaches only 16 % of its volume. The phase shift of ventricular CSF oscillations observed in the numerical results is in good agreement with clinical data. Balédent [4] found a significant phase shift between aqueductal and cervical CSF flow and Wagshul, Eide, and Madsen [60] reported a delay of 15% of the cardiac cycle in the cerebral aqueduct, which matches the 12% delay between peak aqueductal flow and peak blood inflow in our results. Note that we emphasize a comparison between computational and clinical flow rates and volumes rather than CSF velocities as velocities are highly sensitive to geometrical features.

Balédent [4] observed the reversal of cervical CSF flow at the brain equilibrium phase at approximately 55 % of the cardiac cycle. In contrast, in our numerical results, the flow reverses after 38 % of the cycle, which corresponds to the begin of net blood outflow of the cranium. Additionally, their outflow curves take a smooth single-peaked shape over the cardiac cycle, while our results indicate a close resemblance of the flow rate curve and the blood inflow curve. This discrepancy may be explained by a lack of sufficient compliance in the modeled cranial system, which leads to a direct transfer of blood inflow to cervical CSF outflow morphology. Similarly, the multiple reversals of ventricular flow in our model do not match the clinically observed, almost sinusoidal waveforms [4]. These flow reversals are also expected to reduce the corresponding stroke volume. This behaviour might be explained by deviating elastic properties of the brain tissue, leading to multiple oscillations of pressure and flow after the initial excitation of the system at peak blood inflow.

Our model predicts peak ISF velocity magnitudes in agreement with reported values for interstitial bulk flow on the order of micrometers per second [38]. However, the ISF flow computed is pulsatile in time (representing back-and-forth motion over the cardiac cycle rather than bulk flow), and its spatial average is more than two orders of magnitude smaller than its peak value.

The magnitude and direction of the displacement are in good agreement with clinical findings. Based on MRI techniques, Enzmann and Pelc [21], Greitz et al. [24] and Poncelet et al. [42] reported the peak displacement of brain tissue to range from 0.1 to 0.5 mm. More recently, Pahlavian et al. [40] found a peak mean displacement of the brain’s substructures of up to 0.187 *±* 0.05 mm and Sloots, Biessels, and Zwanenburg [48] reported peak displacements of around 0.2 mm; both fit well with the maximal value of 0.22 mm observed in our study. Both these studies [40, 48] reported largest displacements at the brain stem, aligning well with observations from our model. Greitz et al. [24] found a funnel-shaped movement in the dorsal direction and hypothesized that the relatively low pressure below the foramen magnum during early systole induces this motion, which aligns with our numerical results.

Although the model of intracranial pulsatility developed in this work is highly detailed in terms of geometry and biophysical mechanisms, several limitations remain. First, the complex interplay of arterial blood inflow, intracranial dynamics, and venous outflow is simplified into a spatially uniform fluid source in the parenchymal tissue. While the equivalence of the effect of additional fluid volume justifies this approximation, it may still be necessary to include heterogeneities in the source term and differentiate between fluid networks to account for differences in blood perfusion in different regions. Furthermore, even though the time series of net blood flow used in this study (from Balédent [4]) is representative for healthy adults, individual differences in shape and amplitude of the cerebral blood inflow might have a substantial influence on flow and pressure patterns. We also neglect CSF production effects, here without loss of relevance, as any net flow of CSF from its sites of production to absorption is two orders of magnitude smaller than the cardiac induced pulsatile motion [59].

Additional limitations include the uncertainty associated with material parameters, and the assumption of spatial homogeneity in brain tissue, as white and gray matter and subregions likely possess different elastic properties [11]. We expect the effect of moderate heterogeneity on the computational quantities of interest to be relatively small in light of our results with increased elastic stiffness (model B). Furthermore, the boundary conditions describing the transition to the spinal compartment are based on simplifying assumptions. Incorporating a flow resistance to the spinal outflow boundary condition and relaxing the no-displacement assumption of the spinal cord are likely to affect the computational predictions, especially in the brain stem and spinal compartment, and also the pulsatile flow patterns in the aqueduct. Despite the high degree of spatial detail of our model, some features of the intracranial anatomy remain unresolved. As an example, we hypothesize that the tentorium cerebelli would stabilize the brain tissue and block CSF flow, potentially leading to higher pressure differences between the infratentorial and supratentorial regions of the brain.

## Conclusion

In summary, we have presented a new computational model of intracranial fluid flow and tissue motion during the cardiac cycle that offers high resolution and detail in both space and time, and is well-aligned with clinical observations. The model offers a qualitative and quantitative platform for detailed investigation of coupled intracranial dynamics and interplay, both under physiological and pathophysiological conditions.

## Ethics approval and consent to participate

Not applicable.

## Consent for publication

Not applicable.

## Availability of data and materials

The code used to generate and analyze the datasets during the current study are openly available in repository [13].

## Author contributions

M.C., V.V, and M.E.R. designed the computational model and conceived the numerical experiments. M.C. segmented and meshed the MR images. M.C. implemented the simulation algorithms and conducted the simulations. M.C., V.V, and M.E.R. analyzed and discussed the results. M.C. made the figures. M.C, V.V. and M.E.R. wrote the manuscript. All authors revised and approved the final manuscript.

## Supplementary Material

### Mesh & time convergence

A highly detailed mesh is required to adequately resolve the intricate geometry of the human brain and its environment. To ensure a sufficiently fine mesh, we uniformly refined our mesh twice and hence obtained three meshes with increasing resolution (coarse, mid and fine) (Figure 8). Computing the set of quantities of interest on all meshes reveals that the temporal pressure variation in the lateral ventricles and the spinal stroke volume do not change with the mesh resolution, while the aqueductal stroke volume and the peak aqueduct flow rate increase from the coarse to the mid resolution meshes, but remain almost constant in the next refinement stage. The peak displacement and peak transmantle pressure gradient exhibit small decreases from the mid to fine resolution meshes, indicating that further mesh refinement may be desirable. However, given the small changes and limited computational resources, we consider the numerical error acceptable. Similarly, we conduct a time step refinement study on the fine resolution mesh, computing the quantities of interest using 80, 160 and 320 time steps per cardiac cycle. While the temporal pressure variations in the lateral ventricles and the spinal stroke volume again stay constant over time step refinement, the aqueduct stroke volume, the peak aqueduct flow rate, the peak displacement and the peak transmantle pressure gradient slightly increase with the number of time steps.

**Figure 8.**
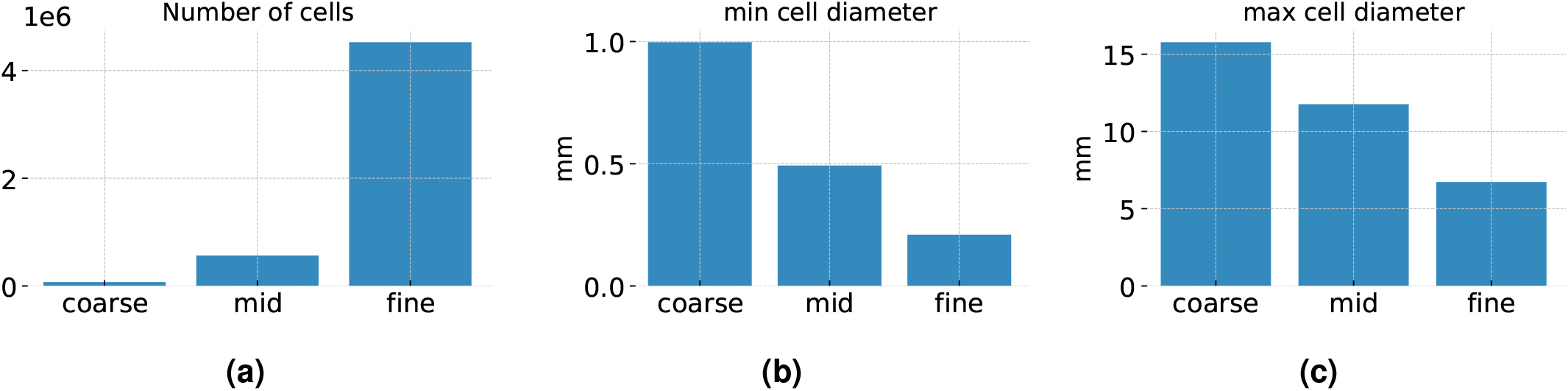
**a)** number of cells of the fine, mid and coarse resolution meshes generated by uniform refinement); **b)** minimal cell diameter of the meshes; **c)** maximal cell diameter of the meshes

**Figure 9.**
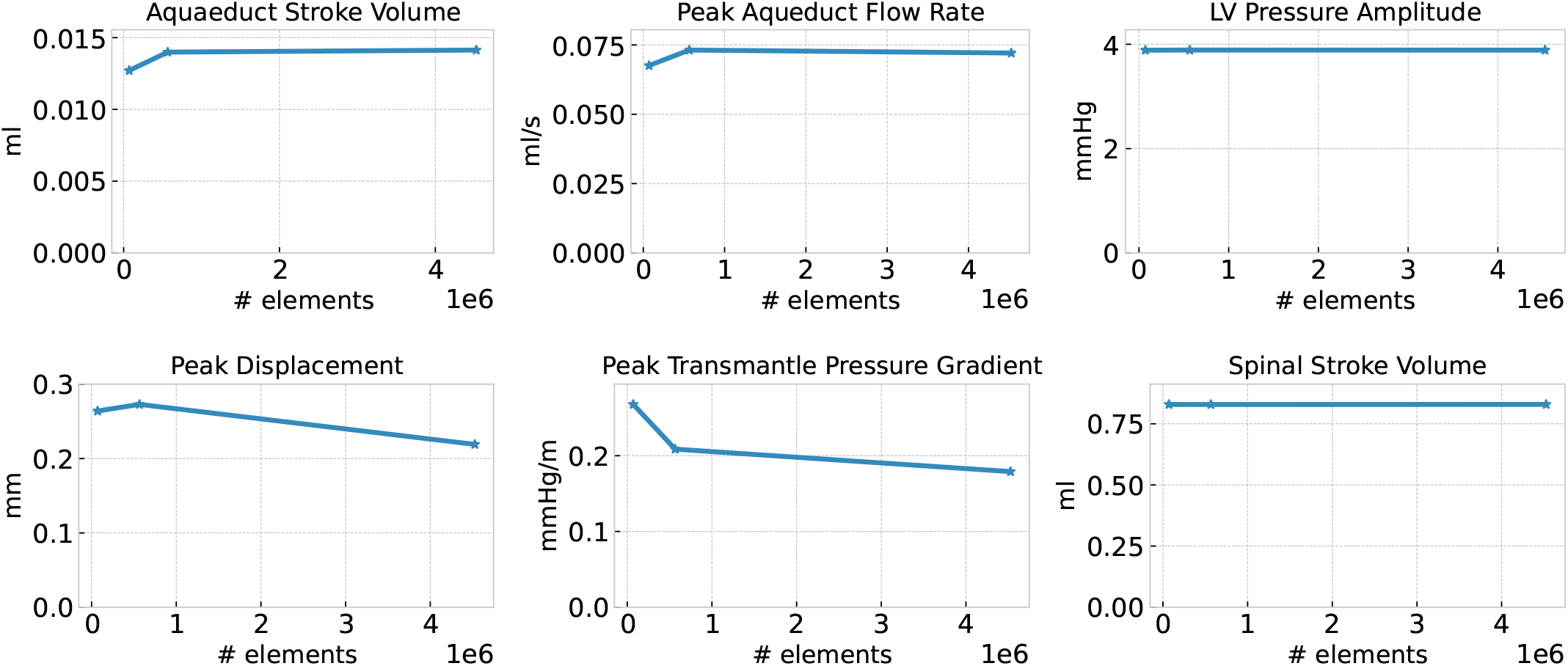
Quantities of interest computed for a sequence of uniformly refined meshes (coarse, mid, fine) with a fixed number of time steps (320 time steps per cardiac cycle).

**Figure 10.**
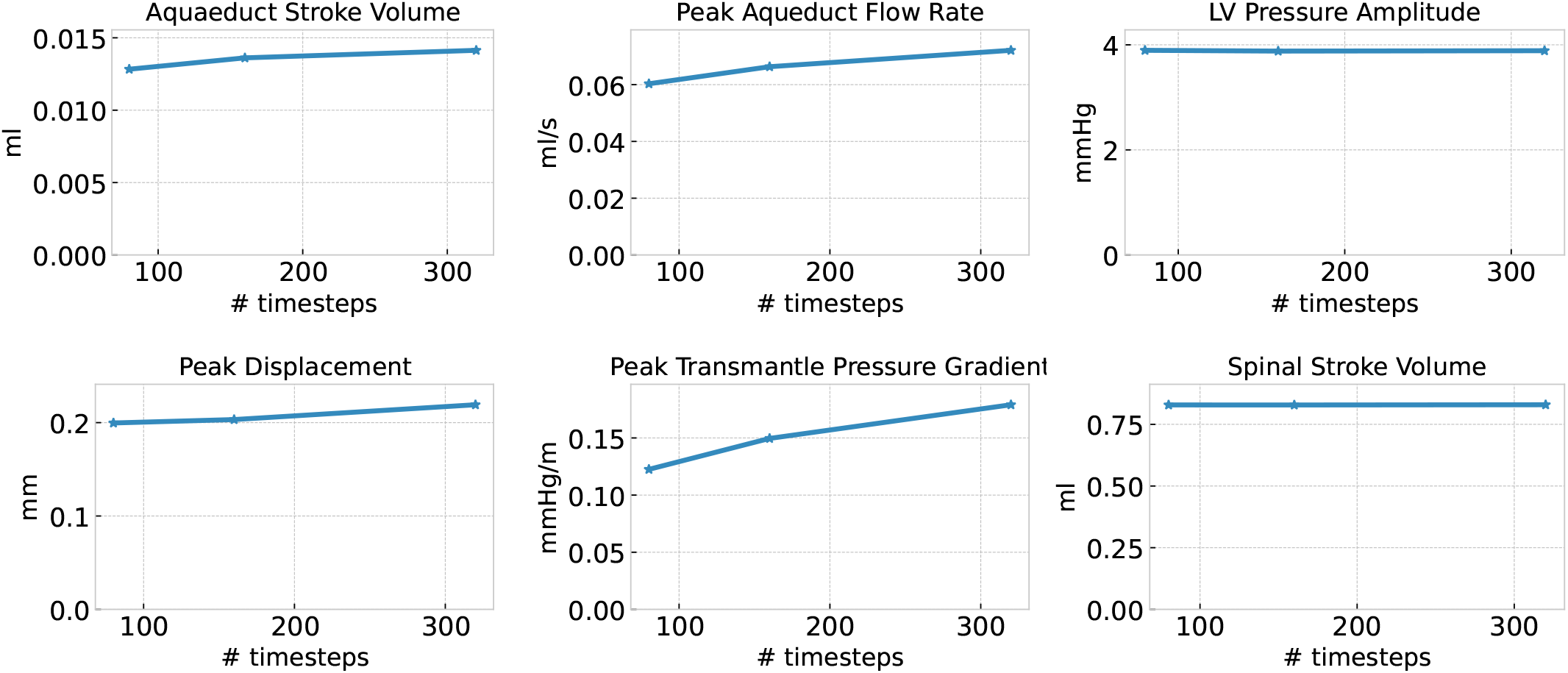
Quantities of interest computed for different numbers of time steps per cardiac cycle (80, 160, 320) on the fine mesh (uniformly refined twice).

### Intracranial Pressure with increased spinal compliance

The ICP pressure curve of Model A (increased spinal compliance) shows a smaller nadir-to-peak amplitude compared to the standard model, but features multiple peaks (P1, P2, P3) per cardiac cycle (Figure 11).

**Figure 11.**
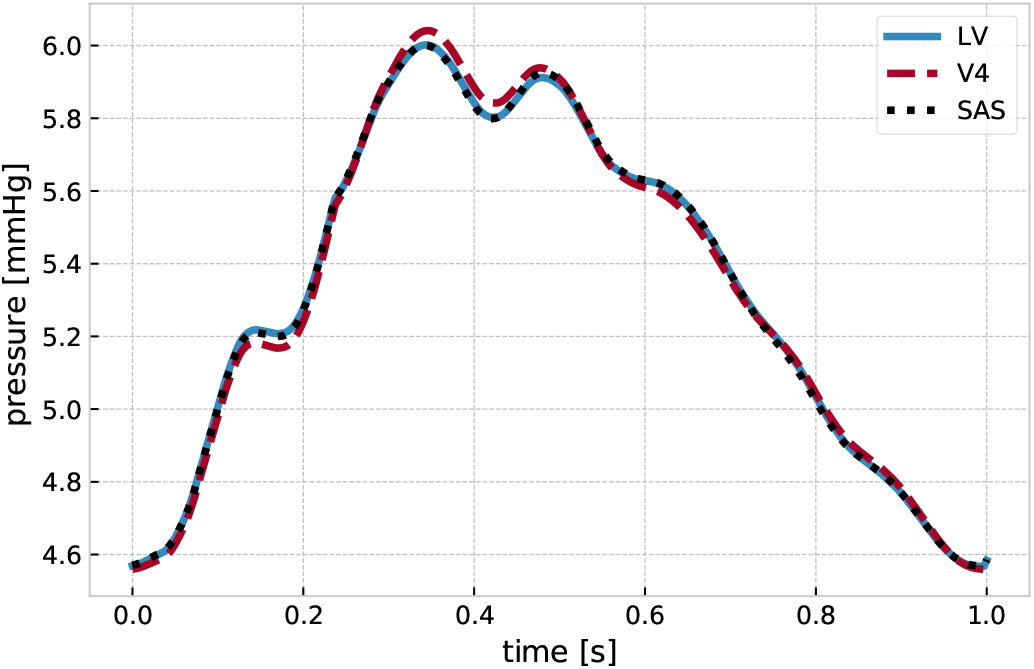
The ICP pressure curve in the lateral ventricles (LV) the fourth ventricle (V4) and the SAS of Model A (increased spinal compliance) shows multiple peaks per cardiac cycle (P1, P2, P3).

